# Discovering Patterns in Bipartite Networks

**DOI:** 10.1101/2022.07.16.500294

**Authors:** Luciano da Fontoura Costa

## Abstract

Bipartite networks constitute an especially important type of network that can be widely applied to model and study several intricate systems in many scientific areas, being also theoretically related to several network models and concepts. However, given a bipartite network the identification of specific recurrent patterns of interest often represents a substantial challenge. In the present work, we apply the coincidence methodology for translating data into networks as a means for automatically identifying repeating interconnection patterns in given bipartite networks. The important issue of normalization of the links strength is also addressed. Though the method is illustrated with respect to foodweb networks, its application is general and encompasses many other subjects and areas.

## 1 Introduction

Bipartite networks constitute a specific type of network or graph characterized by having its nodes subdivided into two main groups *A* and *B*. Typically, interconnections are allowed only between nodes belonging to each of these two sets. As such, bipartite networks focus on *binary* or pair-wise relationships between two types of entities, such as customer/product, consumer-producer, Despite its seeming specificity, bipartite networks are actually found in a large number of problems and areas, from biology to social networks. Some examples include food webs, customer preferences, memberships, and social relationships. Of special importance is that bipartite networks also underly many other types of graphs, such as trees, several planar graphs. In addition, bipartite networks can be applied to model hypergraphs.

Given the practical and theoretical relevance of bipartite graphs as summarized above, a substantial number of related works have been developed and reported in the literature (e.g. [1, 2, 3, 4]). One subject of special relevance while analyzing bipartite networks concerns the identification of patterns of interrelationships shared between several of the involved nodes, which has motivated the application of several community finding approaches.

The present work sets out at investigating the potential of the recently introduced concept of coincidence similarity [5, 6, 7], as well as the derived coincidence method for representing datasets as networks, as the means for detecting patterns of potential interest regarding the relationship between the two sets involved in bipartite structures. Deriving from the Jaccard similarity index (e.g. [8, 9]), but incorporating information about the relative interiority between the vectors being compared while being capable of dealing with real-valued features, the coincidence similarity index has been verified to present enhanced performance in several problems related to pattern recognition (e.g. [7, 10]), reflecting some specific important properties of this similarity index including performing strict comparisons incorporating relative normalization respectively to the vector magnitudes, and presenting substantial robustness to local perturbations (i.e. changes in isolated or small number of features). In addition, the coincidence similarity is particularly capable of implement highly selective and sensitive comparisons.

We start by briefly revising some elementary bipartite network concepts, and follow by discussing what it is in this work understood as patterns of interest in that type of structures. The important issue of weights normalization is addressed next, and then the coincidence methodology for translating datasets into networks is summarized. The methodology for discovering patterns in bipartite structures is presented next, followed by an application example concerning three foodwebs.

## 2 Bipartite Networks Basic Concepts

A bipartite network is henceforth understood to incorporate a set of nodes *A* = {1, 2, …, *N*_*A*_}, a set of nodes *B* = {1, 2, …, *N*_*B*_}, and edges, links or interconnections between nodes from each of these two sets.

Though often represented graphically as being undirected, each of these links indeed corresponds to a pair of directed edges in opposite directions. Given a directed edge, the node from which it originates is henceforth called an *origin node*, while the node at which the edge terminates is referred to as a *destination node*. Observe that edges may proceed from a node in set *A* to a node in *B* or vice-versa, depending of the our specific interest.

The edges can be *binary* (0 = non-existing, 1 = existing) or *weighted* (numeric value), e.g. corresponding to the number of associations between the respective pair of edges.

The *degree* of a node is defined as being equal to the number of edges attached to that node. The *strength* of a node in *A* or *B* corresponds to sum of the weights of the edges attached to that node.

## 3 Patterns of Interest in Bipartite Networks

In this section we define and discuss what we shall understand as patterns of special interest in bipartite networks. Figure 1(a-c) illustrates one typical related situation. The original bipartite network is shown in (a) is small and with relatively few interconnections, but even so it is already quite challenging to visually identify any specific pattern of interconnections as a consequence of the relatively high number of interconnections. However, this bipartite network does contain at least one repeated pattern of interconnection, as illustrated in (b), characterized by three nodes of a set of nodes *A* sharing similar connections with another set of nodes *B*, suggesting the recurrent pattern illustrated in (c).

**Figure 1:**
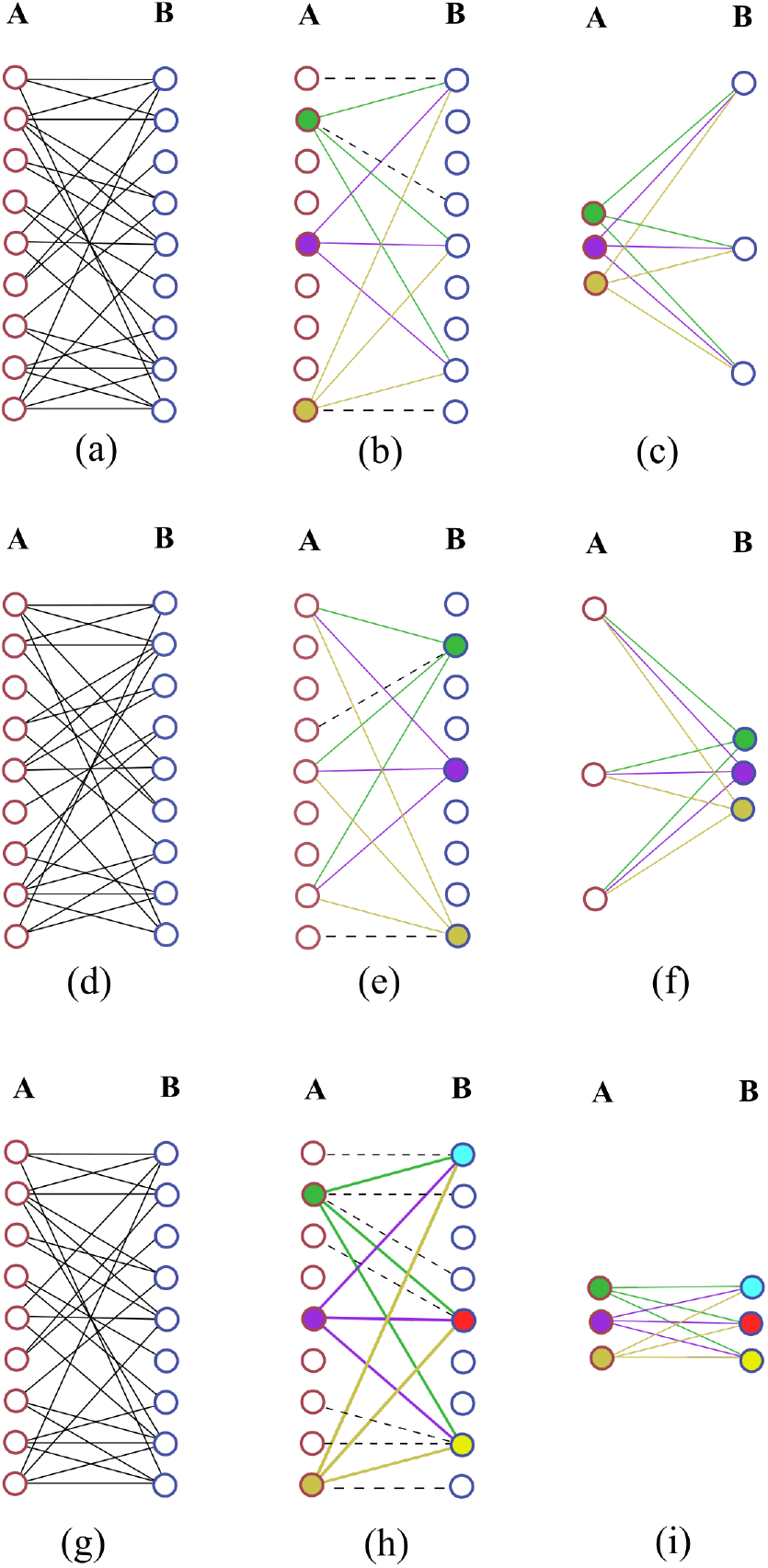
Examples of repeating patterns in a bipartite network. (a-c): an original network (a) in which finding patterns constitute a challenge. One repeating pattern in that network (b), can be summarized as in (c). This pattern refers to the links emanating from network *A*. (d-f): An analogous example concerning the identification of a pattern or interconnectivity with the edges emanating from *B*; and (g-i) Another situation in which the patterns consider the similarity of interconnections emanating from both *A* and *B*.

The key property that is a hallmark of a bipartite pattern is that the destination nodes reached by the respective origin nodes are the same, or *similar* for several origin nodes. For instance, in Figure 1(a-c) we have that the same three destination nodes are reachable from three origin nodes marked as green, violet and yellow.

It is interesting to observe that the pattern in Figure 1(c) has potential interest mainly as a consequence of it recurring three times in this relatively small network. As a matter of fact, the concept of pattern in the area of pattern recognition (e.g. [11, 12, 13]) is strictly associated to groups of instances of patterns that are similar one another while differing from the other available patterns.

A similar situation is illustrated in Figure 1(d-f), but here we are interested in patterns emanating from *B* into *A* in the original bipartite network (d), one of which is depicted in (e), being summarized in (f).

Figure 1(g-i) presents yet another possible situation of interest, in which one is interested in identified common patterns extending from *A* and *B* simultaneously, such as that shown in (h), which is then summarized as (i). Given that the patterns extend toward both sides of the bipartite structure, we suggest them to be called *bippatterns*.

While all situations depicted in Figure 1 involve repetitions of identical patterns, for the sake of enhanced generalization the present work considers a generalization in the sense that small differences between the patterns are allowed. Therefore, some tolerance is need for the quantification of similarity between the patterns which, in the present work, is implemented by adopting the coincidence similarity [5, 6, 7] described in Section 5.

Recurring similar patterns of relationships defined respectively to one of the three situations above are of particular importance because they suggest that these patterns can play a specially relevant role in the respectively studied system. The identification of these types of patterns, therefore, can contribute significantly to a better understanding of the phenomenon or structure represented as a bipartite graph. Before that, however, we briefly discuss the important pre-processing concerning features normalization.

## 4 Weights Normalization

Given a dataset in which each data element is characterized in terms of respective *features*, it constitutes an issue of special relevance whether to normalize or not the feature values, and which normalization to adopt.

The main point here is that distinct features presenting distinct ranges of variation may have respective impact on the results that is proportional to the features with larger magnitudes. In addition, it is often the case that the features have different physical units, which also imply an intrinsic analytical heterogeneity. For these reasons, the procedure known as *standardization* (e.g. [13]) is often adopted, consisting in the application of the following linear statistical transformation to each features *i* for each data element *j*:

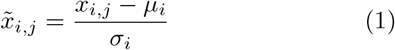

where *µ*_*i*_ and *σ*_*i*_ are the average and standard deviation of the feature *i* along all elements *j*.

In addition to yielding dimensionless new features, standardization also lead to new values with null average and unit standard deviation. In addition, most of the transformed values become comprised in the interval [*−*2, 2]. As with any application involving statistical estimation, it is important that enough data samples are available so as to have reasonable significance. For instance, the standard deviation tends to vary strongly along the first initial samples, so that many observations are often required to achieve satisfactory stability.

Though standardization can often contribute to obtaining more definite results, it should not necessarily applied in every situation. Because standardization removes the average of the set of values of a given feature, (e.g. [14]), + it should not be applied when the average is to be taken into account. In addition, the standardization also normalizes the standard deviation to unit value, implying it not to be adopted when differences of standard deviation magnitude are to be considered. The type of normalization to be applied to a specific dataset depends strongly not only on the characteristics of that dataset, but also on the questions being asked about the data and the type of analysis and methods to be subsequently employed.

While the above considerations relate to general datasets, bipartite networks have some special characteristics that should be considered while normalizing the respective edges weights. Observe that we will be considering these weights as features to be normalized or not. Perhaps the most important aspect consists in the fact that all links in a bipartite network often have the same type (e.g. the number species that are taken by a given consumer). Now, the range of the weight magnitudes of the edges arriving on each specific target node will depend on each specific problem. For instance, we may have that each target node will receive edges with similar weight magnitude ranges, or that quite distinct weight values are observed. Standardization may be considered when too distinct weight magnitudes are involved, provided the average of the weights are not of particular importance for each specific application.

## 5 The Coincidence Methodology

The coincidence similarity index has been introduced [5, 7, 6] as a combination, more specifically the product, of the real-valued Jaccard and interiority (or overlap, e.g. [15]) similarity indices.

Let 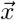 and 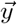 be two real-valued vectors with *N* components each. The *interiority index* between them can be calculated as:

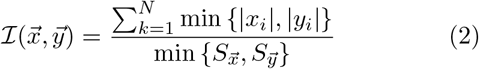

where:

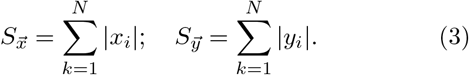

Observe that 0 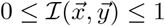.

The real-valued Jaccard index parametrized by *α* between the vectors 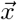 and 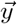 can be expressed as:

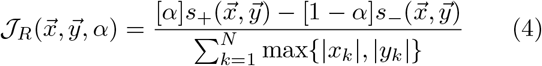

where:

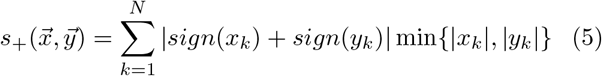

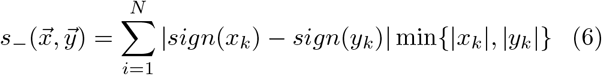

The parameter *α* controls the relative contribution of pairwise values of features with the same or opposite signs. Higher values of *α* will yield larger coincidence similarity values.

The product of Equations 2 and 3 corresponds to the *coindicence index* between the real-valued vectors 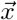 and 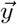. i.e.:

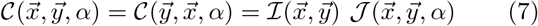

when *α* = 0.5, we have that 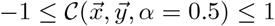.

The coincidence method [16] for translating datasets, where each data element is characterized in terms of *M* features or measurements, involves representing each data element as a node while respective pairwise coincidence values are taken as weights of associated edges. By varying the parameters *α* and *T* it is possible to optimize the obtained networks respectively to several requirements, including maximum modularity (e.g. [17]).

When the coincidence is applied to generate networks as described above, larger values of *α* will yield more connected networks, while larger values of *T* will generate less connected networks.

## 6 Discovering Patterns

Figure 2 illustrates the basic scheme adopted for translating the bipartite table (a) into a respective coincidence network (b) regarding the situation in which the relationships proceed from nodes in *A* into nodes in *B*. Each of the *N*_*A*_ nodes *A*_*i*_ of *A* are represented as a respective node. The weights of the edges of the interconnections between two nodes *A*_*i*_ and *A*_*j*_ of *A* are then determined by the respective coincidence similarity between the feature (weight) vectors *w*(*A*_*i*_, *B*_*k*_) and *w*(*A*_*j*_, *B*_*k*_) in the bipartite table, with *k* = 1, 2, …, *N*_*B*_.

**Figure 2:**
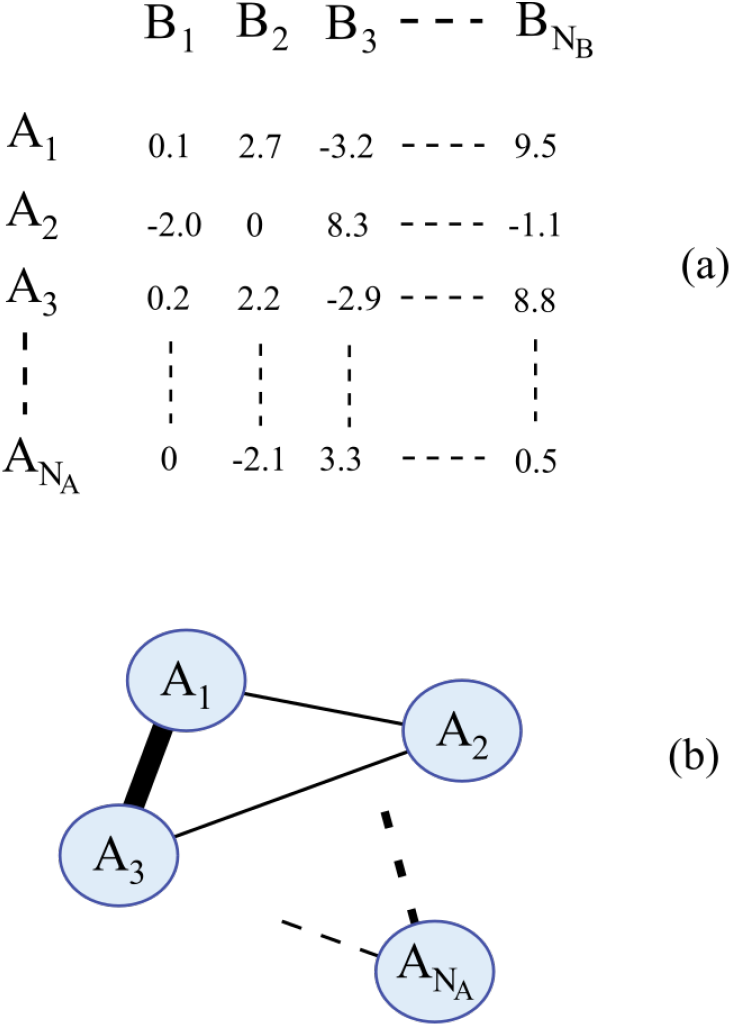
Basic scheme for transforming a bipartite table (a) into a respective coincidence network (b) while considering edges emanating from *A* to *B*. Each entry in *A* is mapped into a respective coincidence network node. The links between these nodes are assigned weights corresponding to the coincidence similarity between the respective features (the connections to the nodes in *B*). The thicker link between *A*_1_ and *A*_3_ reflects the greater similarity between the respective weights observed in the bipartite table for these two entries.

The method to discover patterns in bipartite networks suggested in the present work consists of implementing the following stages:

1. Take each origin node as a data element characterized by the weight of its connections with respective destination nodes, which are taken as respective feature vectors;
2. Apply the coincidence methodology to the so-obtained dataset, therefore yielding a respective network for each specific configuration of the parameters *α* and *T* ;
3. A configuration is chosen so that clusters can be plainly identified (large modularity);
4. For each cluster of interest, the bipartite links originating at the nodes of that cluster are selected as possible instances of a pattern of connections with similar destination nodes, being possibly highlighted in the bipartite representation (red links in the examples in this work).

Observe that each node in the coincidence network therefore corresponds to one of the origin nodes in the bipartite network of interest, while the connections indicate their respective pairwise coincidence similarity.

In addition to identifying patterns of potential interest in bipartite networks, the proposed methodology also provides an informative description of the bipartite interconnectivity in terms of the respectively derived networks, which can provide additional insights about the respectively represented structure. For instance, the weights of the links in the obtained coincidence networks correspond to the similarity between the possible patterns. It is also possible to consider the strength of the origin nodes bipartite network as the diameter of the coincidence network nodes, therefore reflecting this potentially interesting property in the representation.

## 7 Application Example: Foodwebs

In order to illustrate the methodology described in the present work respectively to a specific problem of interest involving real-world data, in this section we provide some application examples respectively to foodwebs, more specifically the datasets *inouye1988* [18], *bezerra2009* [19], and *kevan1970* [20], available in the *Bipartite R Package* [21]. We have adopted *T* = *α* = 0.3 in all subsequent results.

The original bipartite tables are shown as respective images in Figure 3.

**Figure 3:**
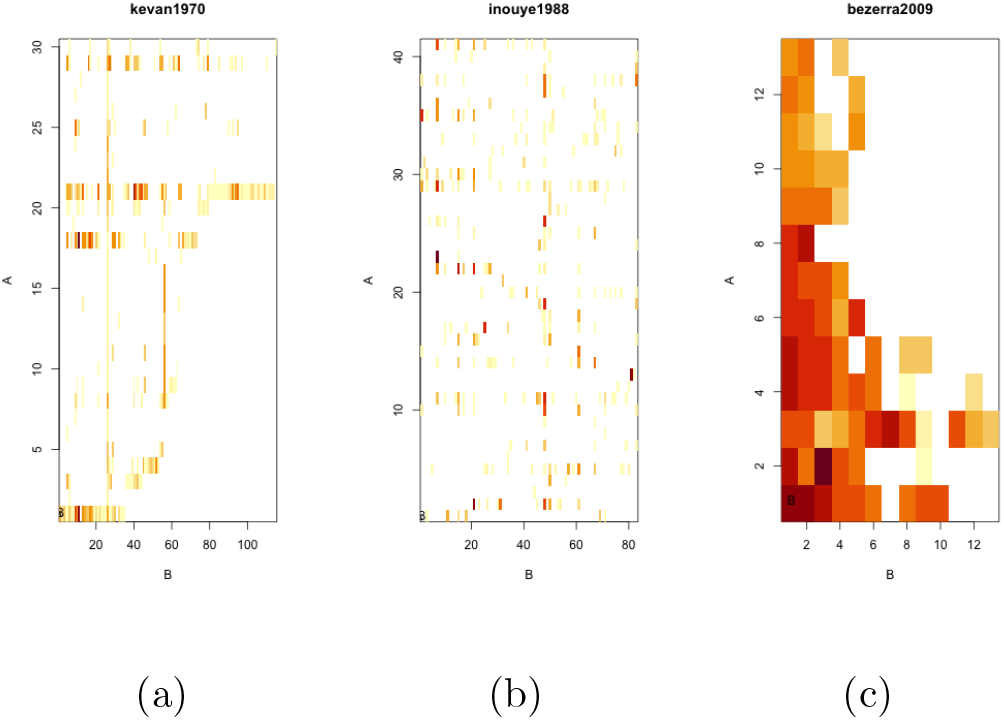
The three original bipartite tables of the three considered foodwebsz shown as respective images. The logarithm of the original weights (number of species interactions) are shown for the sake of a better visualization.

Because the two first datasets involve substantial variations of the interaction magnitude values, the weights are taken as corresponding to these values are thresholded at 0. In the case of the third dataset, which involves more commensurate values, the interaction values are taken as respective weights. In all subsequent visualizations of the bipartite networks, the width of the edges reflect the original interactions, being normalized in non-linear manner so as to avoid large variations in the case of the two first datasets.

Figure 4 presents the coincidence graph(a), coincidence matrix (b) and respective bipartite network (c) obtained from the *kevan1970* dataset [20]. In this particular case, the presence of a well-defined cluster could be already predicted from the block to be found in the coincidence matrix. However, even so it is by no means straightforward to visually identify the pattern of relationship respectively taking place in the respective bipartite structure.

**Figure 4:**
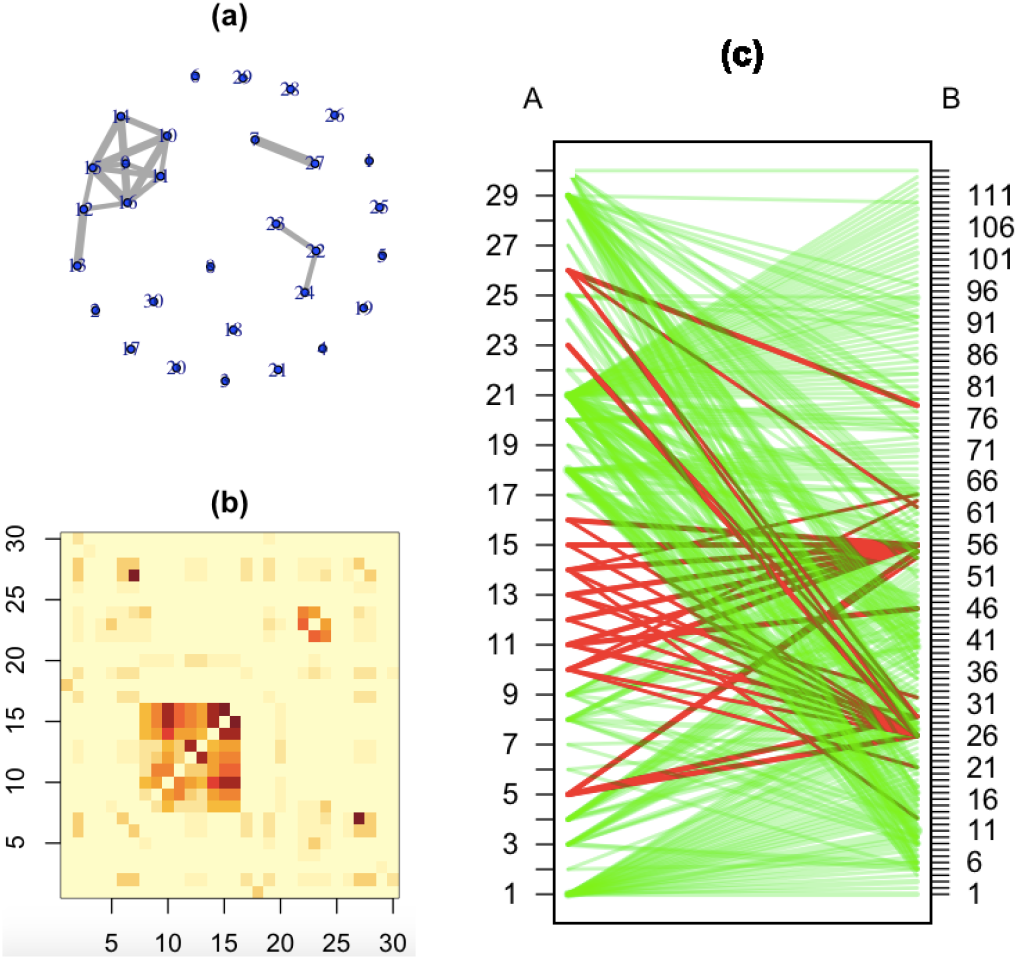
The coincidence graph (a), obtained for *α* = *T* = 0.3, coincidence matrix (b), and bipartite network obtained for the *kevan1970* dataset [20]. The edges corresponding to the largest cluster in (a), taking the nodes in *A* as origin of the edges, are highlighted in red.

The largest module obtained in the coincidence matrix, including nodes from 9 to 16 defines the pattern of potential interest shown in (c). Observe that this pattern is characterized by all origin nodes (9 1o 16) sharing similar destination nodes, which can be identified by the two convergences of edges at the right-hand side of Figure 4. This can also be understood as the identified origin nodes in *A* sharing relationships with the nodes in *B*. Observe some pattern statistical variance reflected in some isolated destination nodes, which were generalized by the adopted parameters. Two other small modules can be observed in Figure 4(a) which could be further investigated.

The application of the suggested methodology yielded the bipartite dataset *inouye1988* [18] shown in Figure 5. Because the larger variation of counts in this dataset cannot be suitably standardized given its sparsity, the counts were thresholded by 1. We can observe that three entries in *A* present similar relationships with nodes in *B*, especially 48 and 83.

**Figure 5:**
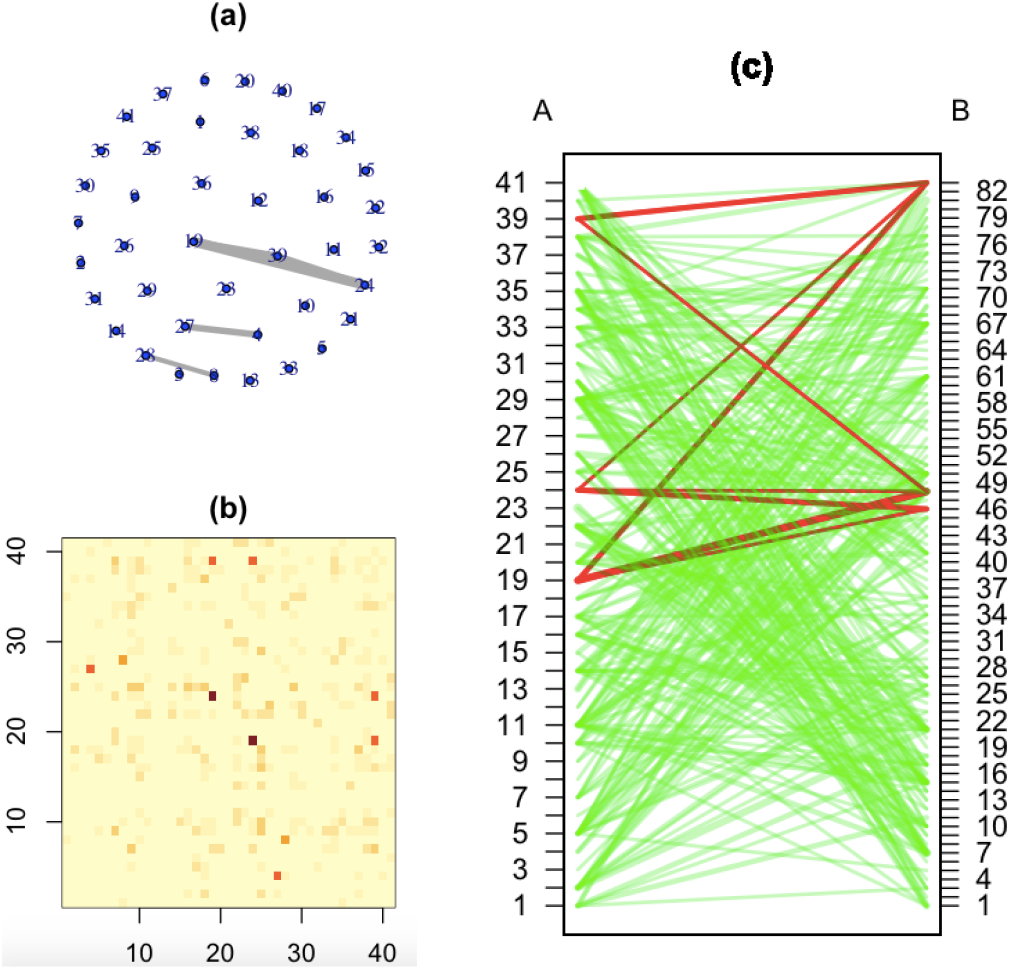
The coincidence graph (a), obtained for *α* = *T* = 0.3, coincidence matrix (b), and bipartite network obtained for the *inouye1988* dataset [18]. The edges corresponding to the largest cluster in (a), taking the nodes in *A* as origin of the edges, are highlighted in red

Figure 6 illustrates the results obtained for the dataset *bezerra2009* [19]. Again, little insights can be gathered from direct visualization of the original bipartite or even the the obtained coincidence matrix. The application of the coincidence methodology led to the identification of a respective network containing a dominant cluster composed of 7 nodes which, when highlighted in the respective bipartite visualization, indicates a pattern of particulr interest corresponding to the fact that two nodes in *A* mostly relate to four similar destinations in *B*, namely node 1 to 4 in that set, allowing for some relatively small fluctuations in the respective interconnecting patterns. It is also interesting to observe that the detected cluster can be understood to present two modules, one corresponding to nodes 4 to 6 in *A*, which are more interconnected with the nodes 1 to 4 in *B*, as well as the smaller group defined by the nodes 9, 10 and 13 in *A*, which also share connections with the other group, but in a slightly distinct and/or less connected manner (recall that the weights of the interactions are taken into account by the coincidence).

**Figure 6:**
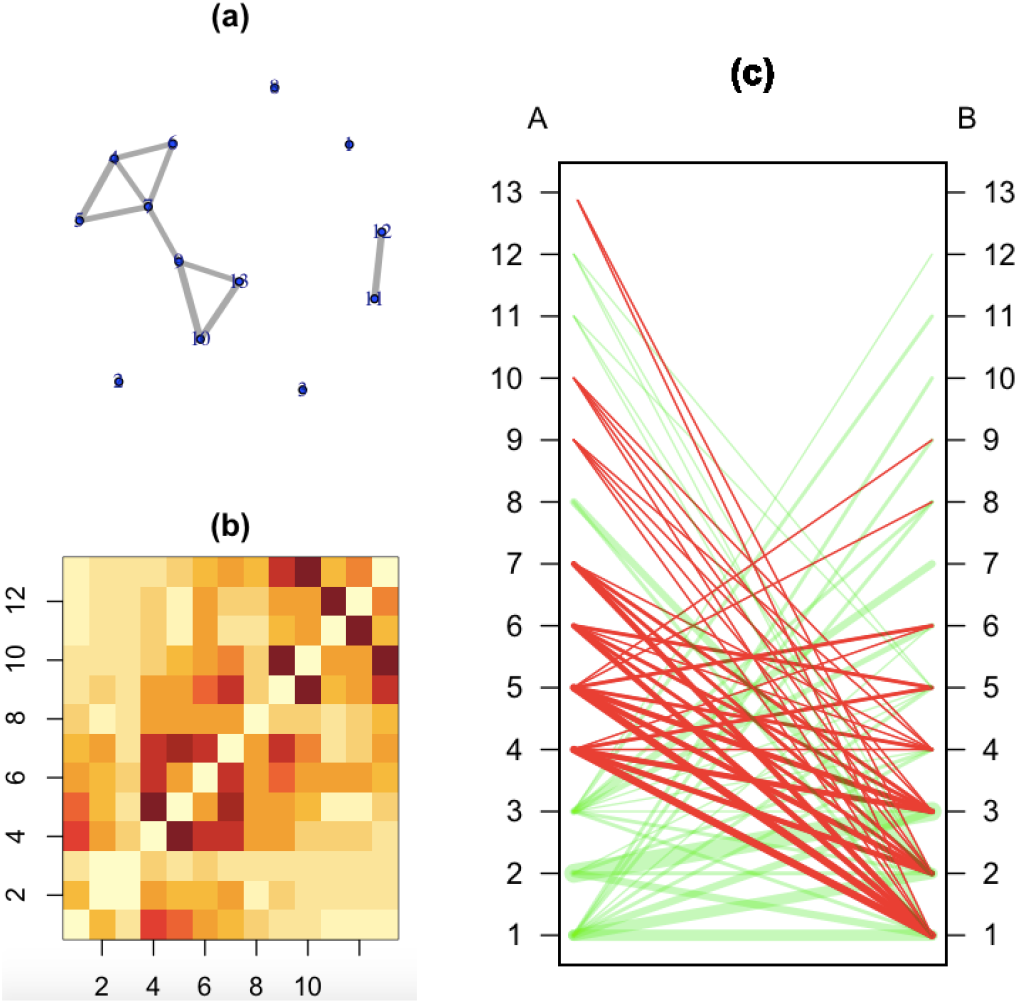
The coincidence graph (a), obtained for *α* = *T* = 0.3, coincidence matrix (b), and bipartite network obtained for the *bezerra2009* dataset [19]. The edges corresponding to the largest cluster in (a), taking the nodes in *A* as origin of the edges, are highlighted in red. A relatively large module (or community) could be identified from the obtained coincidence network.

## 8 Concluding Remarks

Despite their seeming specificity, bipartite networks can be actually applied to a wide range or problems in many scientific areas, while also theoretically underlying several other types of networks and mathematical structures.

One important problem while analyzing bipartite networks concerns the identification of repeating patterns of specific interest, such as several links being established between similar sets of nodes at both sides of a bipartite structure. While this task can hardly be performed through human visual inspection, automated methods need to be developed for that finality. The present work described one possible approach, based on the concept of similarity between the patterns of interconnections as gauged by the coincidence similarity [5, 6, 16].

The potential of the reported approach has been illustrated respectively to three real-world foodweb datasets, allowing the identification of respective patterns of interest related to the bipartite interconnectivity that could be of intrinsic interest in the context of related domains. The identification of significant modules (or communities) in the obtained coincidence networks led to the immediate identification of particularly interesting patterns of interrelationships between the nodes at both side of the bipartite networks in all the considered situations.

Several further developments are motivated by the reported concepts, methods and results, including applications to several other areas, automatization of the suggested procedure by monitoring the coincidence network modularity (e.g. seeking for maximum modularity by using [17]), as well as construction of networks of interrelationships between the identified motifs by using the own coincidence methodology.

## Acknowledgments

Luciano da F. Costa thanks CNPq (grant no. 307085/2018-0) and FAPESP (grant 15/22308-2).

